# A Scalable Distributed-Memory MPI Implementation of Smith-Waterman with Token-Passing Traceback

**DOI:** 10.64898/2026.09.08.750277

**Authors:** Maryam Fatima, Waqas Ali

## Abstract

As genomic sequencing produces increasingly massive datasets, accurate local sequence alignment via the Smith-Waterman(SW) algorithm remains computationally prohibitive due to its space and quadratic time complexity *O*(*mn*). While parallelization addresses a path forward, existing MPI-based solutions present a critical bottleneck during the traceback phase, either omitting it entirely or gathering the entire direction matrix to a single root node, which severely limits scalability. To overcome this, we propose a fully distributed Message Passing Interface (MPI) implementation featuring a token passing traceback scheme. Our approach distributes the directional matrix across all participating ranks, reducing per-rank memory footprint from *O*(*mn*) to *O*(*mn*/*p*), thereby enabling the alignment of sequences far beyond the capacity of sequential or centralized parallel methods. We validate our method on real human DNA sequences (BRCA1, BRCA2, Titin, chr1, chr2) and synthetic datasets up to 50k×50k. Results demonstrate that this MPI implementation aligns 98k×98k real DNA (chr1×chr2) in 58.9160 seconds across 16 MPI processes, a task where the sequential baseline fails due to out-of-memory errors. We achieve best speedups on synthetic data of 19.30× (40k×40k, 16 MPI processes) and on real DNA data is 13.39× (BRCA1×Titin, 8 MPI processes), while capping per-rank memory for the largest dataset at just 573 MB. By enabling exact, memory-scalable alignment with fully distributed traceback on standard CPU MPI clusters, this work fills a critical gap in high-performance computational genomics.

## 1. Introduction

Bioinformatics is based on sequence alignment. It is important to compare DNA, RNA, and protein sequences to find evolutionary relationships, disease-causing mutations and protein structures. There are two types of alignment – global (Needleman-Wunsch)[1] and local (Smith-Waterman)[2]. Local alignment is more useful since in biological sequences similarity can occur only in small parts of the sequence. This is why local alignment, particularly SW alignment, is more useful and correct for real biological applications.

In 1981, Temple F. Smith and Michael S. Waterman proposed SW algorithm [2], which is regarded as a gold standard algorithm for local alignment since it can provide the optimal local alignment. However, it takes *O*(*mn*) time and space if it is formulated with dynamic programming, where *m* and *n* are the lengths of the two sequences. This means that doubling the number of sequences will quadruple the computation time and memory requirements will be quadrupled. Next-generation sequencing has accelerated genomic data growth substantially [3]. Developing new applications for sequencing data generates terabytes of data that are produced on a regular basis; thus, the need to make the SW algorithm faster and more memory efficient is essential. Without the ability to deal with complexity, genomic analysis on a large scale becomes impractical. Heuristic aligners such as BLAST [4], BWA [5], and Bowtie 2 [6] trade optimality for speed on large databases.

The SW algorithm is broken down into two steps: (1) matrix filling (score computation) and (2) traceback (alignment reconstruction). The similarity score is calculated between the sequences during the filling of the matrix, and during the traceback phase, the actual alignment is reconstructed. Affine gap extensions build on the classical formulation [7]. There are several ways to extensively parallelize matrix filling, such as SIMD, OpenMP, MPI, GPUs, and FPGA [8, 9, 10, 11, 12, 13, 14]. Despite this, traceback is still a bottleneck that is not widely recognized.

Traceback constructs the optimal alignment from the highest scoring cell to the zero cell following the “maximum scores path”. This is a sequential process as each step determines the next. To obtain the direction matrix, *O*(*mn*) memory is required to store direction matrix for accurate traceback. If we also store the full score matrix, then the total memory will be 2 × *O*(*mn*), which is not feasible for long sequences. Existing parallel implementations either skip traceback (return only maximum score) or perform it serially on CPU, but require collecting all matrix data at the root. This is the approach is used by FPGA, GPU, and MPI_Gather designs, and leads to a memory-bottleneck at the root node.

Prior surveys such as Xia et al. [15] emphasize parallel matrix fill, data layouts, and throughput (GCUPS) but do not systematically address distributed traceback or memory-efficient traceback on CPU MPI clusters. Bonab et al. [16] report Actor-Serial memory usage of 746.92 GB on the Titin dataset at 40 cores, while traceback is coordinated by a central manager actor. In manager-based approaches, partial alignment data are consolidated centrally, which can become a communication and memory bottleneck. The SWIMM 2.0 implementation [17] does not implement traceback and focuses on score-only computation with AVX-512. LMSWA [18] is a fully serial memory-optimized variant. In these ways, traceback remains either absent, centralized, or serial.

Our work addresses this need by implementing distributed-memory MPI with rank-local direction storage and token-passing traceback. We have four innovations in our approach. First, the direction matrix is decomposed to store it across MPI ranks. The memory per node decreases from *O*(*mn*) to *O*(*mn*/*p*) with number of ranks, *p*. Large sequences may be processed without running into memory problems. Second, traceback without gathering the full direction matrix or using a central manager is performed by using a token-passing approach. The propagation of the token (current column index and completion status) happens up the ranks. Each rank applies segment to its local direction matrix, and passes the token to the next rank. Using this method, the full matrix does not need to be stored on a single node. Third, the score matrix is calculated by using the rolling buffers. The only block that stores DP matrix is the current block, which significantly saves memory. Last, the output is correct with respect to the sequential baseline. Each alignment is verified against the sequential baseline.

Experimental results demonstrate that our approach can align 98k×98k real DNA (chr1×chr2) in 58.9160 seconds on 16 MPI processes with 573 MB memory per rank, while the sequential baseline fails for this size (OOM). The best speedup was obtained on real DNA dataset (BRCA1×Titin) which is a factor of 13.39×, and on synthetic dataset of 40k×40k it is 19.30× using 96.72 MB memory per rank at 16 MPI processes.

The remainder of the paper is laid out as follows. Section 2 (Related Works) provides background on the SW algorithm, its computational problems and the previous parallel SW implementations. Section 3 (Sequential Baseline) presents the sequential baseline pseudo-code, complexity, and execution times. Section 4 (Proposed Algorithm Design) discusses our methodology, including MPI pipeline architecture, distributed direction matrix storage and token-passing traceback mechanism. Section 5 (Experimental Results) gives experimental setup and results such as execution time, speedup, memory usage and scalability analysis. Section 6 (Discussion) compares with state-of-the-art implementations found in the literature. Section 7 (Conclusion and Future Work) concludes the paper with future work.

## 2. Related Works

The Smith-Waterman (SW) algorithm has been extensively investigated concerning parallelization on different architectures such as SIMD, OpenMP, MPI, GPUs, FP-GAs, Actor models, and Xeon Phi coprocessors. We give a detailed overview of these implementations, including an analysis of their traceback strategies, to show that there is a key and long-standing need in the literature: a fully distributed traceback mechanism on CPU clusters.

The existing literature addresses the SW traceback phase in one of four main ways. In our suggested approach, we add a new fifth category:

### 2.1. SIMD and Vectorization Implementations

To speed up the matrix filling step of the SW algorithm, the most popular approach is the SIMD (Single Instruction Multiple Data) vectorization, which refers to multiple data streams being processed by a single instruction. Farrar (2007) [8] presented striped layout mapping of query sequences to SIMD registers, which resulted in a 6× speedup over other SIMD implementations on the same hardware. Rognes (2011) [9] presented SWIPE, a system that implemented inter-sequence SIMD parallelization that achieved up to 220 GCUPS. Further developments included the Parasail C library by Daily (2016) [19] and the SeqAn library by Rahn et al. (2018) [20] which added vectorization and multi-threading to reach 194 GCUPS.

These implementations are in Category A. They only perform the database search and do not execute the traceback phase at all, only reporting the highest score.

### 2.2. Shared Memory (OpenMP) Implementations

Khan et al. (2015) [10] used an anti-diagonal wavefront approach with OpenMP [23] and reported a 2.63× speedup over the sequential baseline. Filling the matrix was parallelized, but traceback was performed sequentially on the host after matrix completion.

This is not an MPI_Gather design; matrix fill is parallel, but traceback remains serial on the host.

### 2.3. GPU and Hardware Accelerators

Due to the huge number of cores, GPUs are often used for SW acceleration. Liu et al. (2013) [13] developed CUDASW++ 3.0, achieving 185.6 GCUPS on a single GPU (Category A, score-only). Wang et al. (2023) [12] proposed GPUSW with a 1.66× speedup (without traceback) and 1.53× (with traceback). Hybrid CPU–GPU designs with unified memory have also been explored [24].

GPUSW is in Category C. The GPU computes the direction matrix while the traceback is done serially by one thread of the GPU. Sandes et al. (2014) [22] developed CUDAlign 3.0, but it was score-only and traceback was omitted because the overhead of memory and complexity of multi-GPU coordination were significant.

FPGAs are known to be very energy-efficient for SW acceleration. Nanou (2025) [14] proposed a multi-kernel implementation on the Alveo U250 FPGA, achieving speedups of up to ~14× over a single-thread CPU baseline. Rucci et al. (2018) [25] created an OpenCL-based SWIFOLD (Category A, score-only).

Nanou’s approach is in Category C. The direction matrix is calculated by the FPGA, but traceback reconstruction is performed on the host CPU after FPGA computation, which can create a host-side memory bottleneck for very large matrices.

### 2.4. Distributed Memory (MPI) Implementations

Distributed-memory systems use the standard MPI [26]. Khaled et al. (2016) [11] and Ninama et al. (2025) [21] both proposed hybrid MPI-OpenMP models; Ninama explicitly gathers sub-matrices to the root for serial traceback. Other notable works are those of Zwaka (2024) [27] (CUDA-MPI hybrid for Needleman-Wunsch) and Sêmeler & Dias (2026) [28] (implementing on a cluster of Raspberry Pis, where MPI gave a speedup *<*1×). OpenMPI studies on small clusters report fill-only speedups without traceback [29].

In the examined MPI SW implementations with full trace-back, the direction matrix is gathered at the root, and trace-back is performed serially there (Category B), which causes substantial memory bottlenecks for large matrices.

### 2.5. Manycore and Actor Model Implementations

Rucci et al. (2019) [17] created SWIMM 2.0, which achieved up to 511 GCUPS on Xeon Phi coprocessors and 734 GCUPS using AVX-512 extensions on dual Skylake processors (Category A, no traceback). On the other hand, Bonab et al. (2025) [16] implemented SW-actors in C++ using the Actor Framework (CAF), attaining up to 28× speedup on BRCA1 inter-alignment at 40 cores (the journal article reports up to 22× on 40 cores).

This is a Category D approach. Traceback is coordinated by a central manager actor that gathers partial alignment information. On the Titin dataset at 40 cores, Actor-Serial requires 746.92 GB of memory.

Habeeb et al. (2025) [18] introduced LMSWA, which, by avoiding storage of the full score matrix, reduces runtime memory by up to 88.28% at 20 KB sequence length. The algorithm is entirely serial and extends alignment to sequences up to 110 KB on an 8 GB system, whereas standard SWA fails above 20 KB on the same machine. Theoretically, Wang (2013) [30] has proposed a VLSI architecture that can align in *O*(*m* + *n*) time using *mn* processing elements, but this architecture is not implemented in practice (Category A).

The literature search shows a clear trend: Category A focuses only on the generation of the maximum score (SIMD, Xeon Phi, GPUs). Category B and C exhibit significant memory limitations, such as a serial traceback for a root node, host CPU, or single GPU thread (MPI and OpenMP or FPGA). Category D achieves parallelization but still suffers from unsustainable memory overheads (746 GB on Titin) due to the centralized manager. Xia et al. [15] survey parallel fill strategies and report third-party throughput figures (e.g., SWIPE 220 GCUPS, CUDASW++ 298.8 GCUPS, SeqAn 194 GCUPS), but they do not treat distributed traceback on CPU clusters as a dedicated research direction. We address this gap by distributing the direction matrix and implementing rank-to-rank token-passing traceback without gathering the full direction matrix at one node, thus enabling large-scale distributed sequence alignment in a memory-efficient way.

## 3. Sequential Baseline

This section presents the algorithm of SW sequential baseline 1, its time and space complexity, and execution times obtained on test machine. This baseline acts as the basis for performance evaluation of our MPI implementation. It takes sequences *A* (query, length *N*) and *B* (database, length *M*) as input with scoring scheme match (+3), mismatch (−3), and gap (−2). It returns global maximum score (*S*_max_) and aligned sequences (`alignA`, `alignB`) as output.

There are two important phases, **Matrix Filling:** The score of every cell (*i, j*) depends on its three neighbors (diagonal, up, left) and **Traceback:** Starting from the cell containing maximum score, it reaches zero by following stored directions, which reconstructs the actual alignment. This algorithm computes every cell exactly once. The matrix has (*M* + 1) × (*N* + 1) cells, so time complexity is *O*(*MN*). It stores the full scoring matrix *H* of size (*M* + 1) ×(*N* + 1). With 4-byte integers, total memory is 4 × (*M* + 1) × (*N* + 1) bytes ≈ *O*(*MN*).

### Algorithm 1

Sequential Smith–Waterman Algorithm Baseline

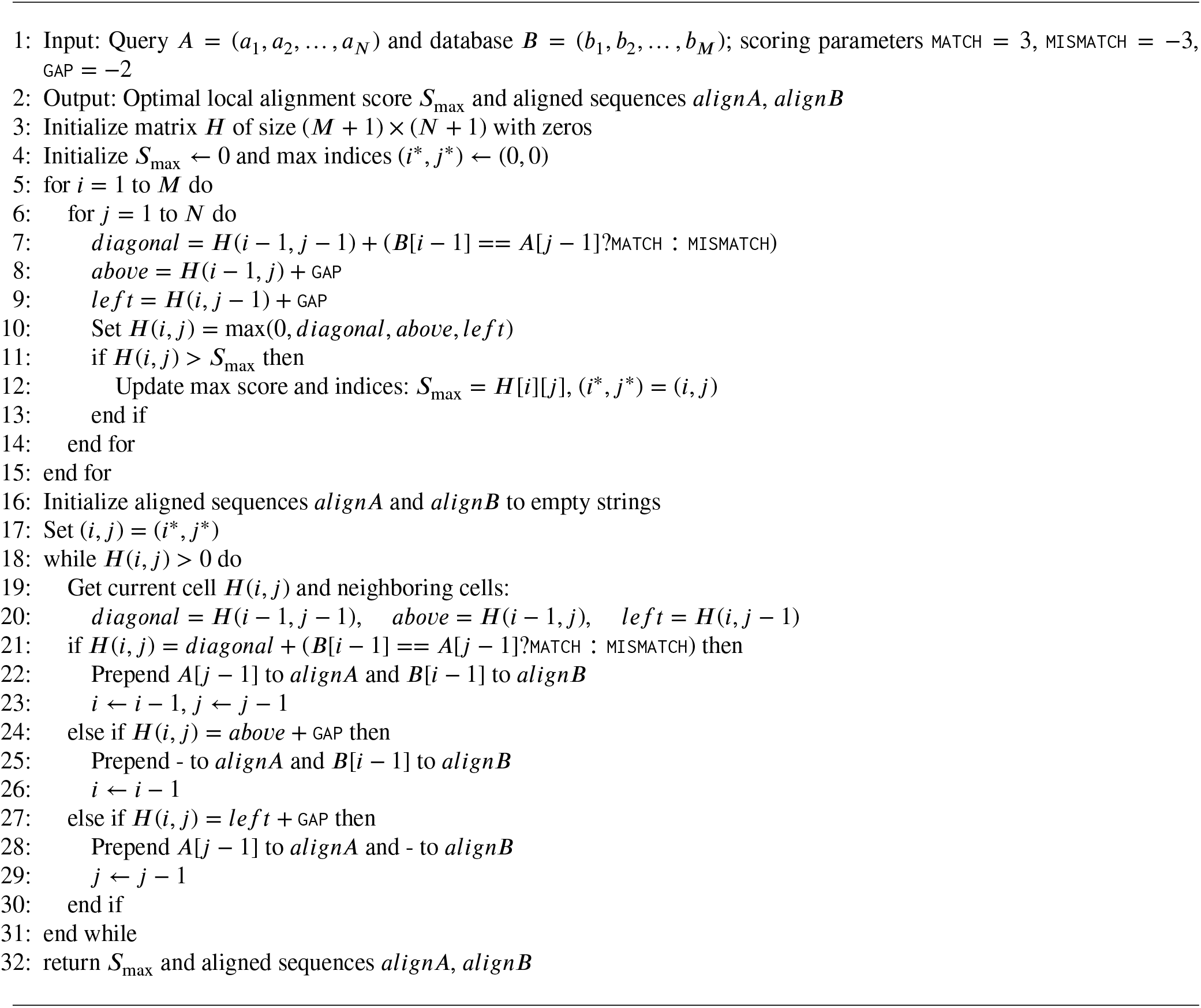

### 3.1. Baseline Execution Times

We tested sequential baseline on synthetic and real datasets. Below are two tables: Table 2 shows execution times on synthetic dataset, whereas Table 3 shows execution times for real datasets.

**Table 1.** Categorization of Traceback Strategies in Literature.

| Category | Description | Examples |
| --- | --- | --- |
| **A. No Traceback** | Does not reconstruct the path of alignment, but simply calculates the maximum possible alignment score. | SWIMM 2.0 [17],<br>CUDASW++ 3.0 [13],<br>SWIPE [9], Parasail [19],<br>SeqAn [20] |
| **B. Gather & Serial (Root)** | Gathers the whole direction matrix to a root node and then performs a serial traceback execution. | MPI_Gather approaches [11, 21] |
| **C. Serial on Host/Thread** | The matrix is computed in parallel, but the serial traceback is done on a single CPU host thread or a single GPU thread. | FPGA [14], GPUSW [12],<br>CUDAlign 3.0 [22] |
| **D. Manager Based (Actor)** | A central actor gathers partial alignments and coordinates the traceback. | SW-actors [16] |
| **E. Distributed (Token-Passing)** | Direction matrix is distributed – traceback is done by passing the token from rank to rank, there is no global gather of the direction matrix. | **Proposed Work** |

**Table 2.**
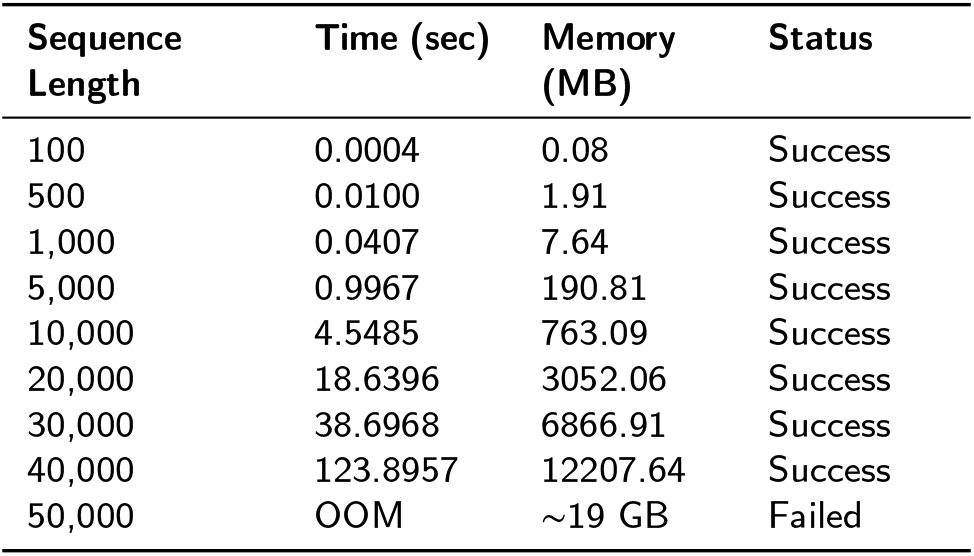
Execution times of sequential baseline on synthetic datasets.

**Table 3.** Execution times of sequential baseline on real gene and DNA datasets.

| Dataset | Sequence Length | Time () | Memory | Status |
| --- | --- | --- | --- | --- |
| BRCA1 × BRCA2 | 7,224 | 3.4199 | 658.99 MB | Success |
| BRCA1 × Titin | 7,224 × 82,029 | 21.0994 | 4521.69 MB | Success |
| Titin × Titin | 82,029 | OOM | ~27 GB | Failed |
| chr1 × chr2 | 98,040 | OOM | ~38 GB | Failed |

Synthetic datasets are generated from random DNA sequences (A, C, G, T). Their purpose is to test scaling behavior in a controlled way. The sequential version executed successfully on datasets up to 40,000×40,000, even though memory usage was 12.2 GB, which exceeds the physical RAM of the test machine (7.7 GB); the OS completed the run using swap memory. For 50,000×50,000, the memory requirement increased to ~19 GB, which was greater than the total available memory (RAM + swap ≈ 9.7 GB), so the process was terminated by the OOM killer.

Real datasets consist of human genes (BRCA1, BRCA2, Titin) and human chromosome fragments (chr1, chr2) from NCBI. These are representative of actual biological applications. BRCA1 × BRCA2 (7k×12k) and BRCA1 × Titin (7k×82k) executed successfully on sequential. BRCA1 × Titin used 4.5 GB of memory, which fit in system memory. Titin × Titin (82k×82k) and chr1×chr2 (98k×98k) both failed because memory requirements (27 GB and 38 GB) exceeded available system memory.

## 4 Proposed Algorithm Design

The proposed algorithm consists of two phases:

In the Forward Pass (Distributed Matrix Scoring) (Algorithm 2), the scoring matrix and direction matrices are computed. The score matrix is computed using rolling buffers (only current block is stored), whereas each rank stores direction matrix for its own local rows.

The second phase is Traceback (Distributed Token-Passing) (Algorithm 3), in which, starting from the cell containing maximum score, a token (containing current column index and completion status) is passed between ranks. Every rank processes segment of its local direction matrix and passes the token to the rank above.

Each rank *P* receives a different portion of a database sequence *T* of length *M*. In this way:

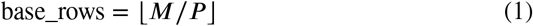

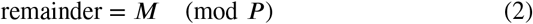

Rank *r* gets these rows:

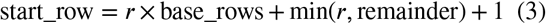

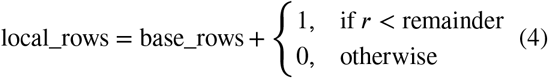

The sequence *S* is split into blocks of fixed size (`BLOCK_SIZE` = 128). All ranks compute all blocks for their local rows, and dependencies (top row, left column, diagonal) are processed through the pipeline. The memory structure per rank is demonstrated in Table 4

**Table 4.** Memory structures maintained per MPI rank.

| Structure | Size | Type | Purpose |
| --- | --- | --- | --- |
| dp_block | $(\text{local\_rows} + 1) \times (\text{BLOCK\_SIZE} + 1)$ | int | Current block score matrix |
| dir_local | $(\text{local\_rows} + 1) \times (N + 1)$ | unsigned char | Direction matrix (1 byte/cell) |
| left_col | $(\text{local\_rows} + 1)$ | int | Previous block last column |
| saved_diag | 1 | int | Diagonal dependency |

The score matrix `dp_block` is stored only for the current block. The direction matrix `dir_local` is stored for the full query length *N*, but only for local rows. In this way, per-rank memory becomes *O*((*M*/*P*) × *N*) bytes per rank, instead of *O*(*M* × *N*). Table 5 shows scoring parameters.

**Table 5.** Scoring parameters used in DPSWA.

| Constant | Value | Description |
| --- | --- | --- |
| MATCH | +3 | Score for nucleotide match |
| MISMATCH | -3 | Penalty for nucleotide mismatch |
| GAP | -2 | Linear gap penalty |
| BLOCK_SIZE | 128 | Columns per pipeline block |

### 4.1. Phase 1: Distributed Matrix Scoring (Forward Pass)

For every block *b*, pipeline communication is managed by the following conditions. If rank *>* 0, then rank *r* receives its top row (`dp_block[0][1..cols]`) from rank *r* − 1 via `MPI_Recv`, `dp_block[0][0] = saved_diag`, and `saved_diag` is updated to `dp_block[0][cols]`. For each local row *i*, `dp_block[i][0] = left_col[i]`. For each cell (*i, j*) local and global parameters are computed and the score matrix is updated:

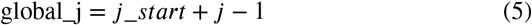

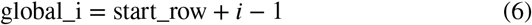

The match score is evaluated on the basis of this condition:

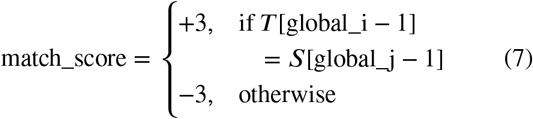

While validating the execution dependencies, cell score is calculated:

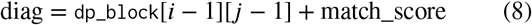

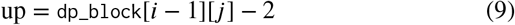

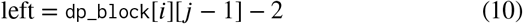

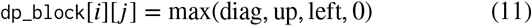

Direction codes for the traceback matrix are given in Table 6. After computing score for each cell, the matching direction is stored by checking backtrack parameters.

**Table 6.** Direction codes for traceback matrix.

| Condition | Direction | Code |
| --- | --- | --- |
| $\text{dp\_block}[i][j] == 0$ | DIR_NONE | 3 |
| $\text{dp\_block}[i][j] == \text{diag}$ | DIR_DIAG | 0 |
| $\text{dp\_block}[i][j] == \text{up}$ | DIR_UP | 1 |
| $\text{dp\_block}[i][j] == \text{left}$ | DIR_LEFT | 2 |

**Table 7.**
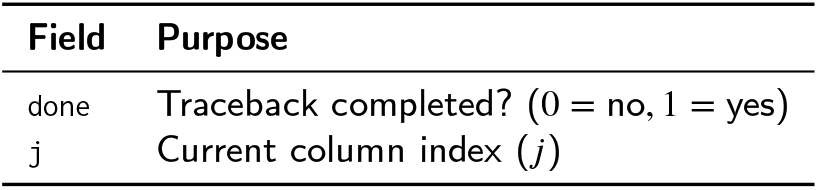
Token fields and their purpose.

| Field | Purpose |
| --- | --- |
| done | Traceback completed? (0 = no, 1 = yes) |
| j | Current column index ($j$) |

The direction matrix is stored in `dir_local`[i][global_j], which is optimized for single column memory footprint. Each rank tracks the local maximum score and its respective global coordinates:

**if** `dp_block`[i][j] *>* local_max **then**

local_max = `dp_block`[*i*][*j*]

max_i_local = global_i

max_j_local = global_j

**end if**

After the block is computed, if rank *< P* − 1, then rank *r* sends its bottom row (`dp_block[local_rows][1..cols]`) to rank *r* + 1 via `MPI_Send` so that the next process can resolve the dependency. All ranks reduce their `local_max` using `MPI_Allreduce` to find the global maximum *S*_max_ and the rank `max_rank` containing it. The `max_rank` then broadcasts global coordinates (global_i, global_j) to all other ranks via `MPI_Bcast`.

#### Algorithm 2

DPSWA Forward Pass — Block-Pipeline Smith-Waterman Scoring

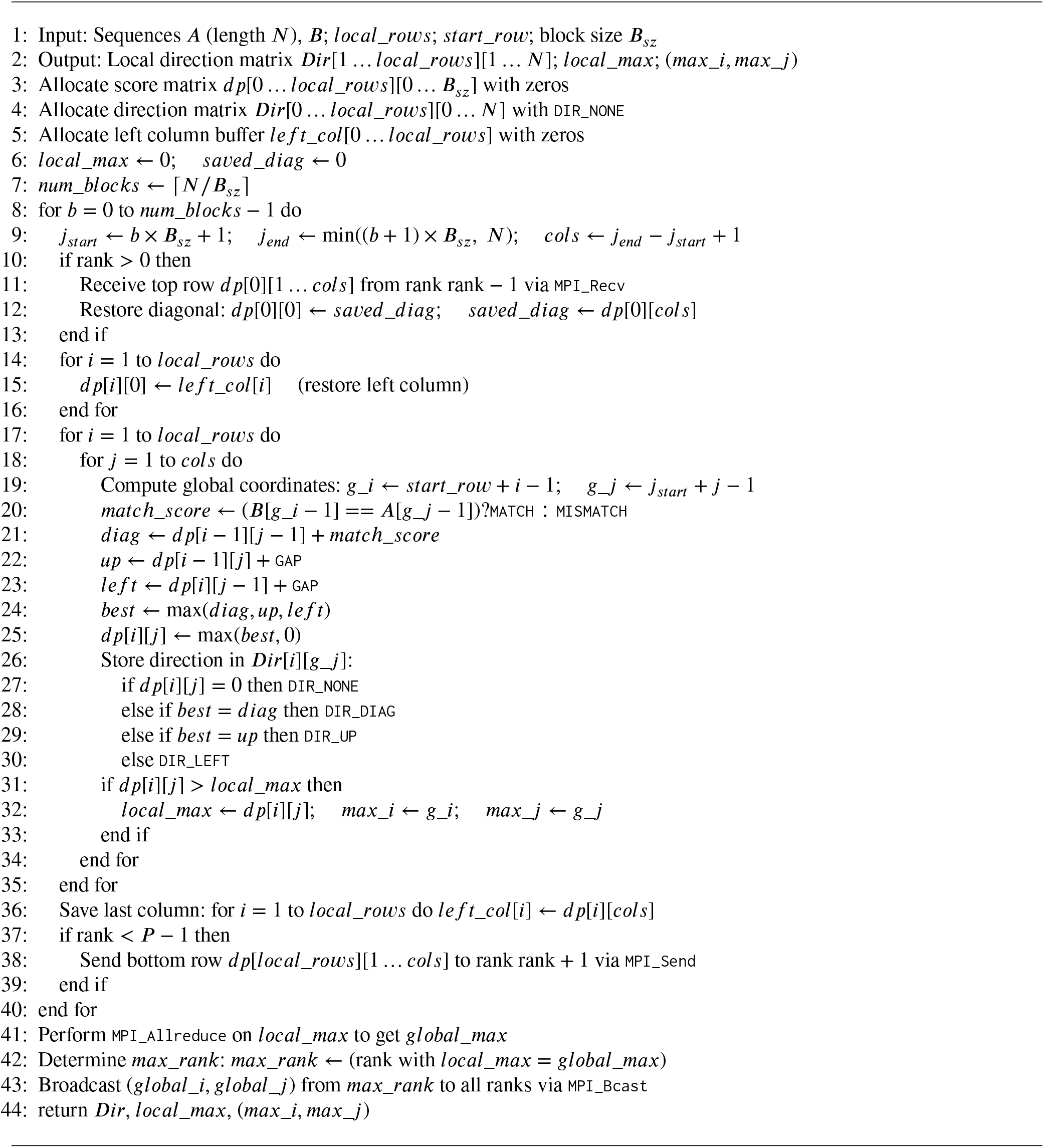

### 4.2. Phase 2: Distributed Token-Passing Traceback

For execution coordination in the traceback phase, a token is used which consists of two primary fields 7.

Token is initialized in a way such that `max_rank` initializes the token: `done` = 0, `j` = global_j. and other ranks wait in the blocking state to receive token from rank *r* + 1. When a rank receives the token, it executes the local loop as per its boundary coordinates:

local_i = global_i − start_row + 1, if rank = max_rank,

= local_rows, otherwise.

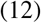

The loop continues until *local*_*i >* 0, *j >* 0 and `dir_local`[local_i][*j*] ≠ `DIR_NONE`. Table 8 shows the traceback directions and corresponding pointer actions. If *local*_*i* == 0 (top boundary reached), the column matrix pointer *j* is updated and the token is passed to the upper rank by setting `done` = 0. Otherwise, `done` = 1 is set, which means the traceback sequence is completed successfully.

**Table 8.** Traceback directions and corresponding pointer actions.

| Direction | Action |
| --- | --- |
| DIR_DIAG | Prepend $S[j - 1]$ to $\text{localA}$, $T[\text{start\_row} + \text{local\_i} - 2]$ to $\text{localB}$, $\text{local\_i} --$, $j --$ |
| DIR_UP | Prepend '-' to $\text{localA}$, $T[\text{start\_row} + \text{local\_i} - 2]$ to $\text{localB}$, $\text{local\_i} --$ |
| DIR_LEFT | Prepend $S[j - 1]$ to $\text{localA}$, '-' to $\text{localB}$, $j --$ |

The token is forwarded on the basis of following conditions: If rank *>* 0, then rank *r* sends updated token (`done`, *j*) to rank *r* − 1 via a point-to-point communication channel using `MPI_Send`. Each rank builds a local alignment fragment (`local_alignA`, `local_alignB`). When processing is complete, Rank 0 gathers all fragments and assembles them into the final global sequence alignments `AlignA` and `AlignB` by concatenation, as shown in Algorithm 4.

### 4.3. Memory Optimization Techniques and Complexity Analysis

In order to reduce memory usage, several techniques have been used, as referenced in Table 9. The first technique is *rolling buffers*, in which the score matrix `dp_block` is stored for the current block only, which eliminates the need to store the full score matrix in memory. This means that the memory requirement can be controlled to an extent, especially in the case of large sequences. The second technique is the *direction matrix-only* approach, where similarity score values are discarded and only the direction matrix is preserved, which takes 1 byte per cell. The third technique is *distributed storage*, in which the direction matrix is stored on each MPI rank for its own local rows, reducing per-rank memory overhead. The last and most important technique is *no global gather*. During traceback the entire direction matrix is not gathered at any single node: instead token-passing scheme is used so that the centralized memory bottleneck can be prevented.

**Table 9.** Memory optimization techniques and descriptions.

| Technique | Description |
| --- | --- |
| Rolling Buffers | The score matrix $dp\_block$ is stored only for the current block ($\mathcal{O}(local\_rows \times BLOCK\_SIZE)$). The full score matrix is not stored. |
| Direction Matrix Only | The similarity matrix (scores) is discarded; only direction matrix (1 byte/cell) is stored. |
| Distributed Storage | The direction matrix is stored on each rank for its local rows only: $\mathcal{O}((M/P) \times N)$ per rank. |
| No Global Gather | For traceback, the direction matrix is not gathered at any single node; a token-passing scheme is used. |

In distributed form, the time complexity of the forward pass is *O*(*N* × *M*/*P*), where the computational burden is divided among *P* MPI ranks. Due to pipeline-based block processing, communication overhead reaches *O*((*N*/BLOCK_SIZE) × *P*). In the traceback phase, the time complexity depends on the path length of the alignment, which is *O*(*L*), where *L* ≤ *M* + *N*, and the communication cost in the worst case is limited to *O*(*P*) token passes. For per-rank memory complexity, the direction matrix is the major component, having size *O*((*M*/*P*) × *N*) bytes, while the DP block and left column are limited to *O*((*M*/*P*) × BLOCK_SIZE) and *O*(*M*/*P*) integers, respectively. Therefore, the total per-rank memory requirement essentially equals *O*((*M*/*P*) × *N*) bytes, as summarized in Table 10.

**Table 10.** Memory complexity components per MPI rank.

| Component | Size |
| --- | --- |
| Direction Matrix | $\mathcal{O}((M/P) \times N)$ bytes |
| DP Block | $\mathcal{O}((M/P) \times BLOCK\_SIZE)$ integers |
| Left Column | $\mathcal{O}(M/P)$ integers |

**Table 11.**
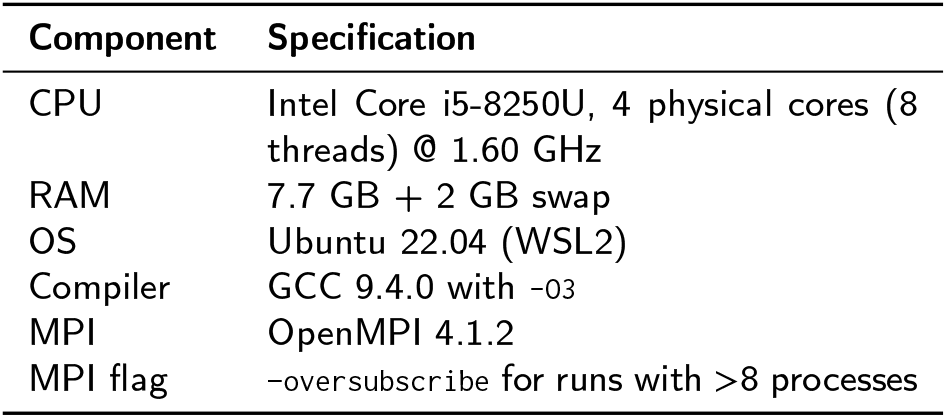
Hardware and software configuration.

**Table 12.** Summary of state-of-the-art parallel SW implementations and their traceback strategies.

| Approach | Key Implementation | Speedup/Perf. | Memory | Traceback Details |
| --- | --- | --- | --- | --- |
| SIMD | Farrar (2007) [8], Rognes (2011) [9] | High Throughput | Full Matrix | Category A |
| OpenMP | Khan et al. (2015) [10] | 2.63× | Full Matrix | Category C |
| GPU | Wang et al. (2023) [12] | 1.53× (with traceback) | Full Matrix | Category C |
| FPGA | Nanou (2025) [14] | ~14× (vs. CPU) | Rolling Buffers | Category C |
| MPI | Khaled (2016) [11], Ninama (2025) [21] | Hybrid MPI–OpenMP | Full Matrix | Category B |
| Xeon Phi | Rucci et al. (2019) [17] | 734 GCUPS | 8-bit biasing | Category A |
| Actor Model | Bonab et al. (2025) [16] | 28× (BRCA1, 40 cores) | 746 GB (Titin, 40 cores) | Category D |
| Mem. Opt. | Habeeb et al. (2025) [18] | N/A | 88% reduction | Serial |
| **MPI** | **Proposed Approach** | **19.30×** | **Distributed** | Category E |

## 5. Experimental Results

This section presents the experimental results of our MPI implementation on synthetic and real datasets. All experiments were conducted on the following platform 11. Results are divided into three categories: Strong Scaling Graphs (Speedup vs. number of processes), Execution Times Tables (complete data for each configuration), and Memory Reduction Analysis (sequential run vs. MPI run).

### Algorithm 3

DPSWA Distributed Traceback — Token-Passing Scheme

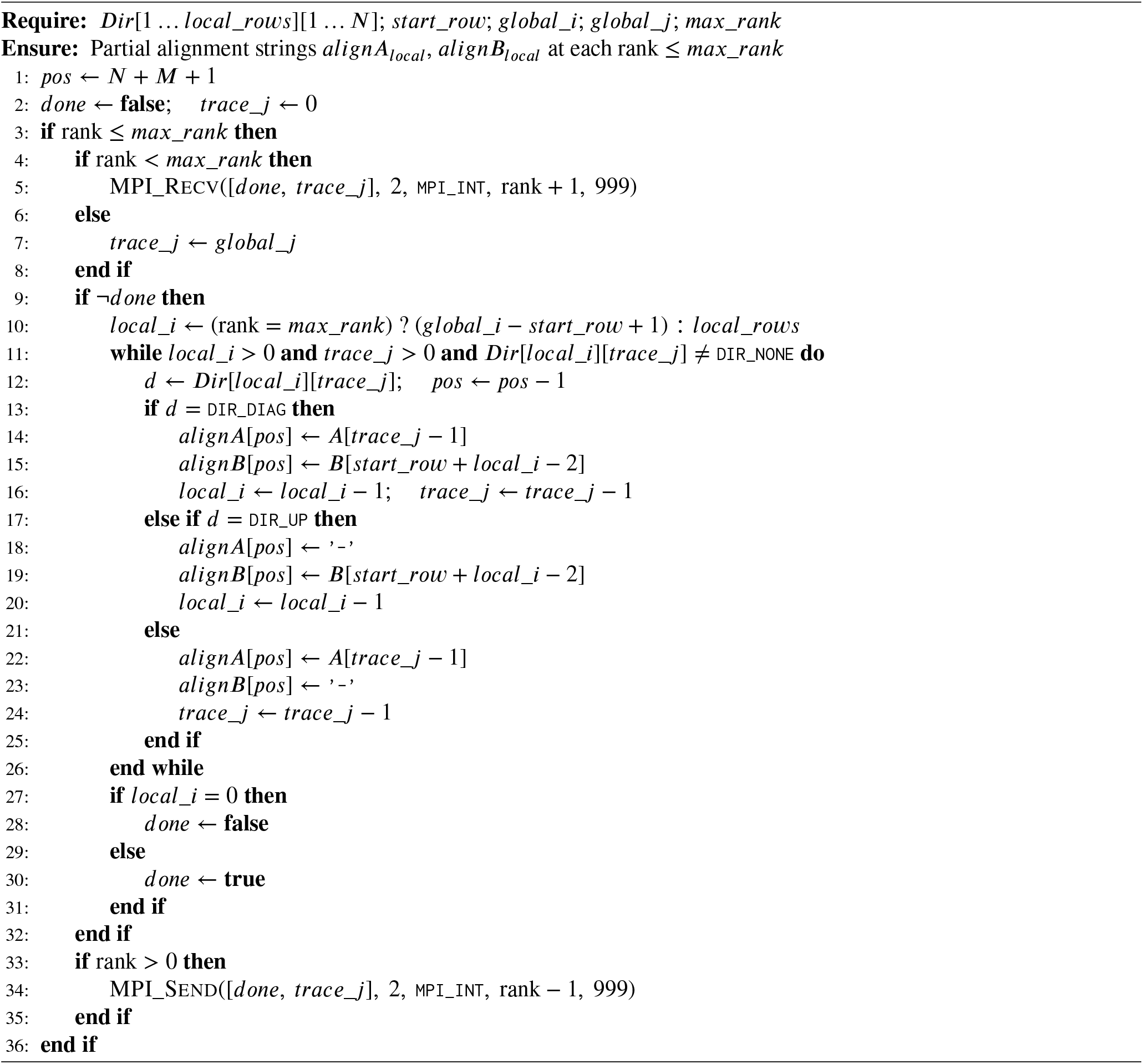

The aim of this analysis is to demostrate that the proposed distributed-memory MPI implementation can process a large sequence efficiently, while the sequential baseline implementation cannot handle such sequences due to Out-Of-Memory (OOM) errors.

### 5.1. Strong Scaling Analysis (Speedup vs Processes)

For these experiments we kept the problem size constant and increased the number of processes to see how well we could take advantage of additional processes with our implementation of MPI.

Figure 1 shows speedup results on synthetic data (5k, 10k, 20k, 30k, 40k, 50k). Initially, speedup increases with the number of processes and reaches at its peak on 8–16 processes, and then declines at 32 processes. This is due to pipeline communication overhead and oversubscription. The maximum speedup (best) is 19.30× at 16 processes.

**Figure 1.**
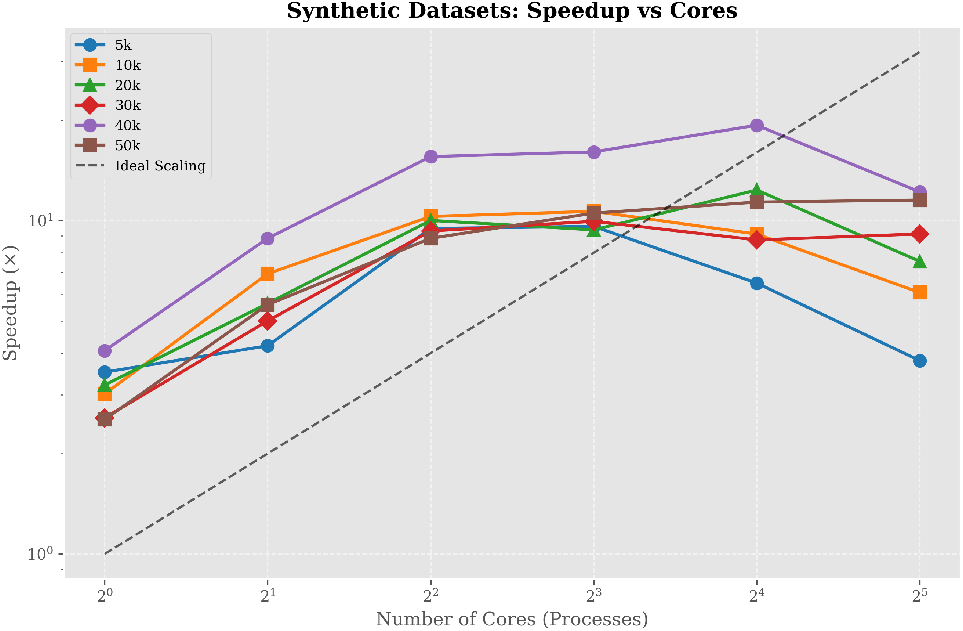
Speedup vs processes for synthetic datasets.

Similarly, Figure 2 depicts the experimental findings with the real datasets. The best speedup for BRCA1×BRCA2 is 12.77× (8 processes) and for BRCA1×Titin is 13.39× (8 processes). These results show that the proposed approach is effective on real biological sequences.

**Figure 2.**
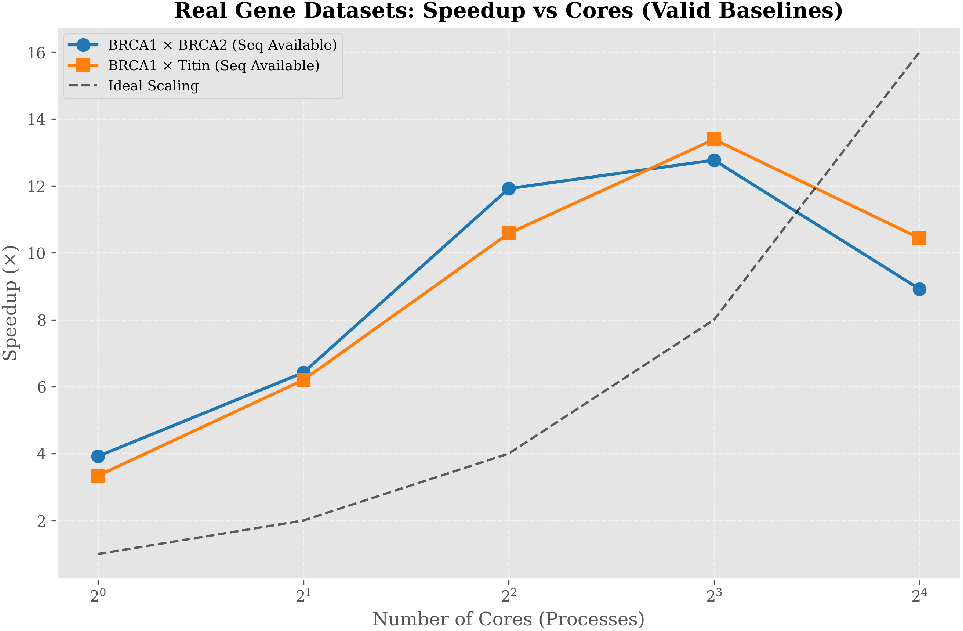
Speedup vs processes for real gene datasets.

The baseline failed in sequential mode, and therefore speedup for Titin×Titin cannot be calculated. Time scaling for Titin is however shown in Figure 3. The 8-core setup acts as the sweet spot, as shown in the strong scaling graphs. Beyond that, the communication overhead becomes the factor that takes over and starts to reduce the speedup, while the memory efficiency remains constant.

**Figure 3.**
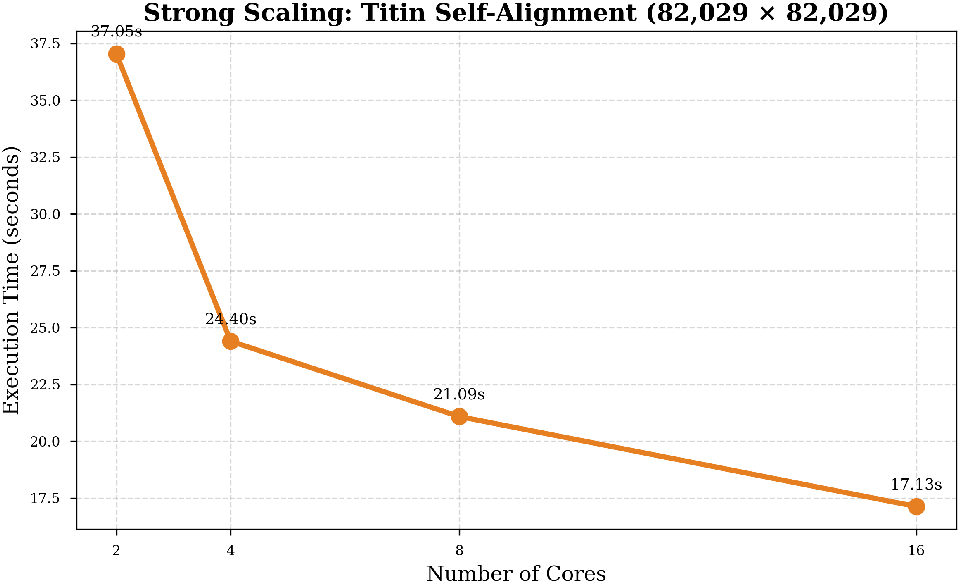
Time vs processes for Titin dataset strong scaling.

**Figure 4.**
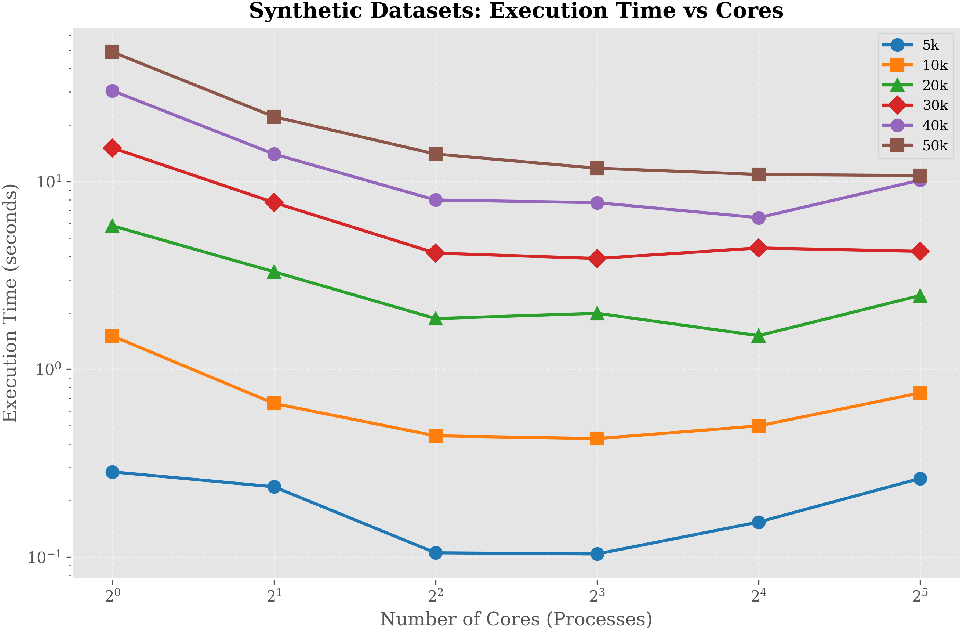
Execution time vs processes for synthetic datasets.

**Figure 5.**
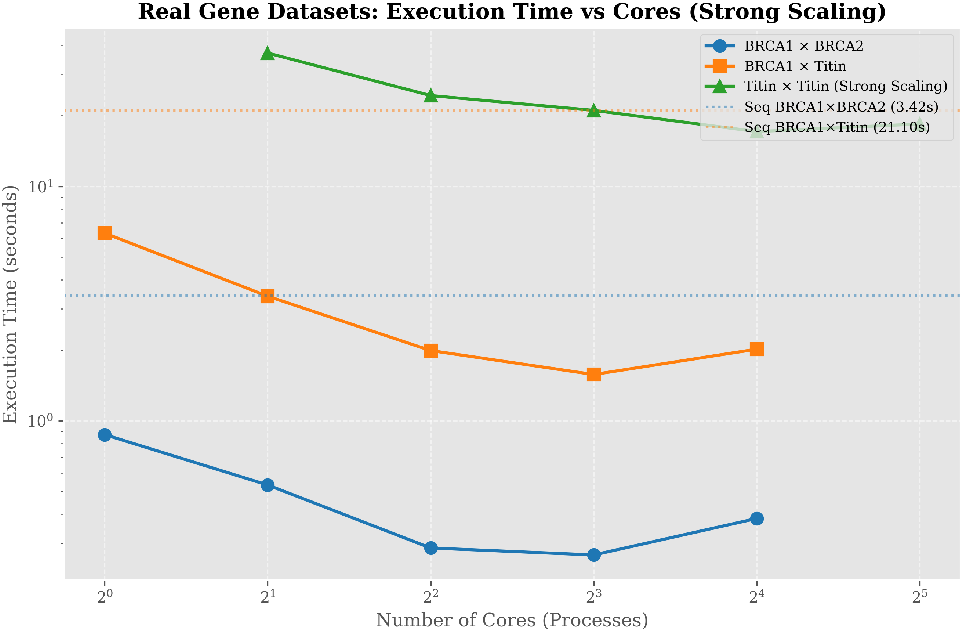
Execution time vs processes for real gene datasets.

**Figure 6.**
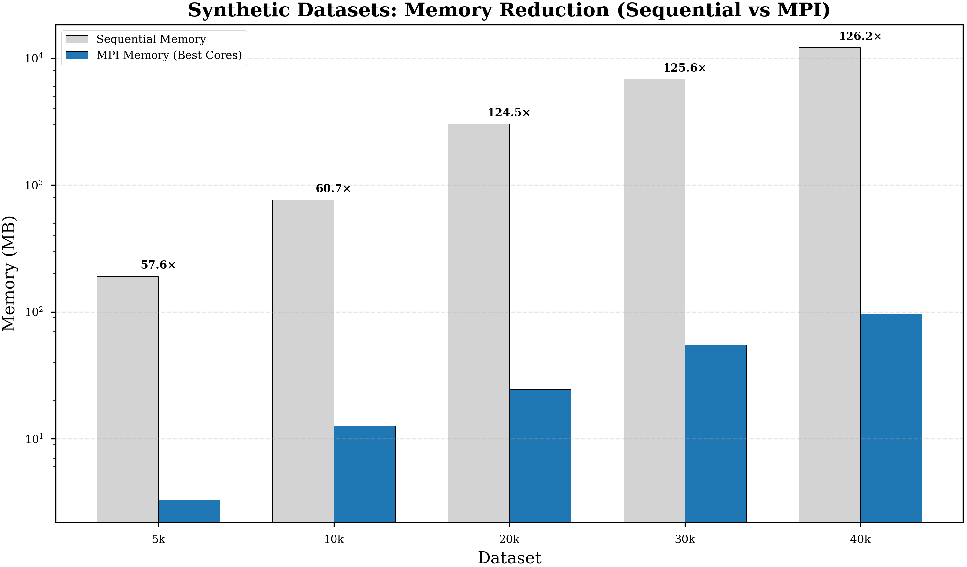
Memory reduction comparison for synthetic datasets.

**Figure 7.**
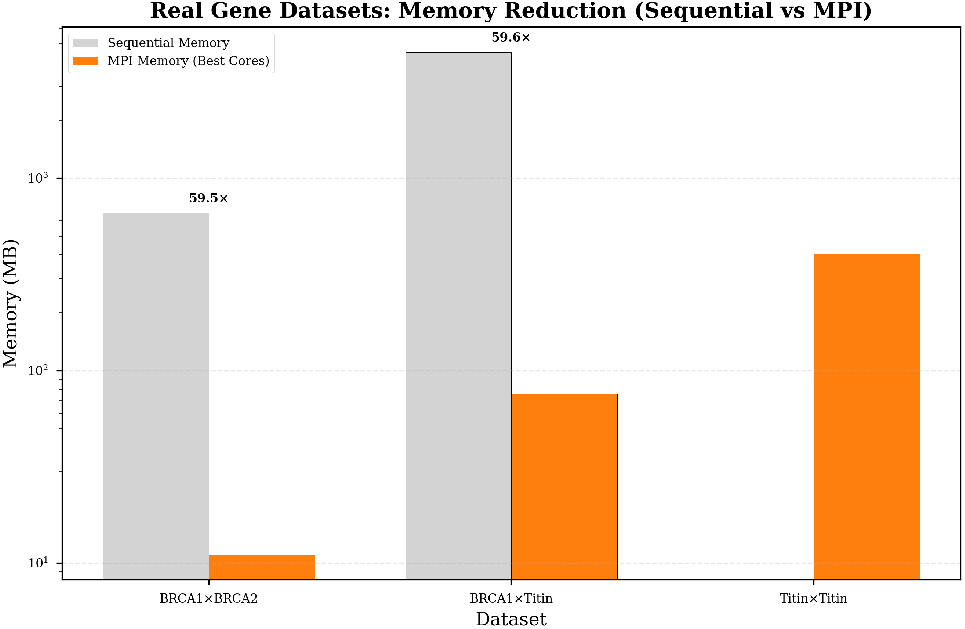
Memory reduction comparison for real gene datasets.

#### Algorithm 4

DPSWA Gather and Output

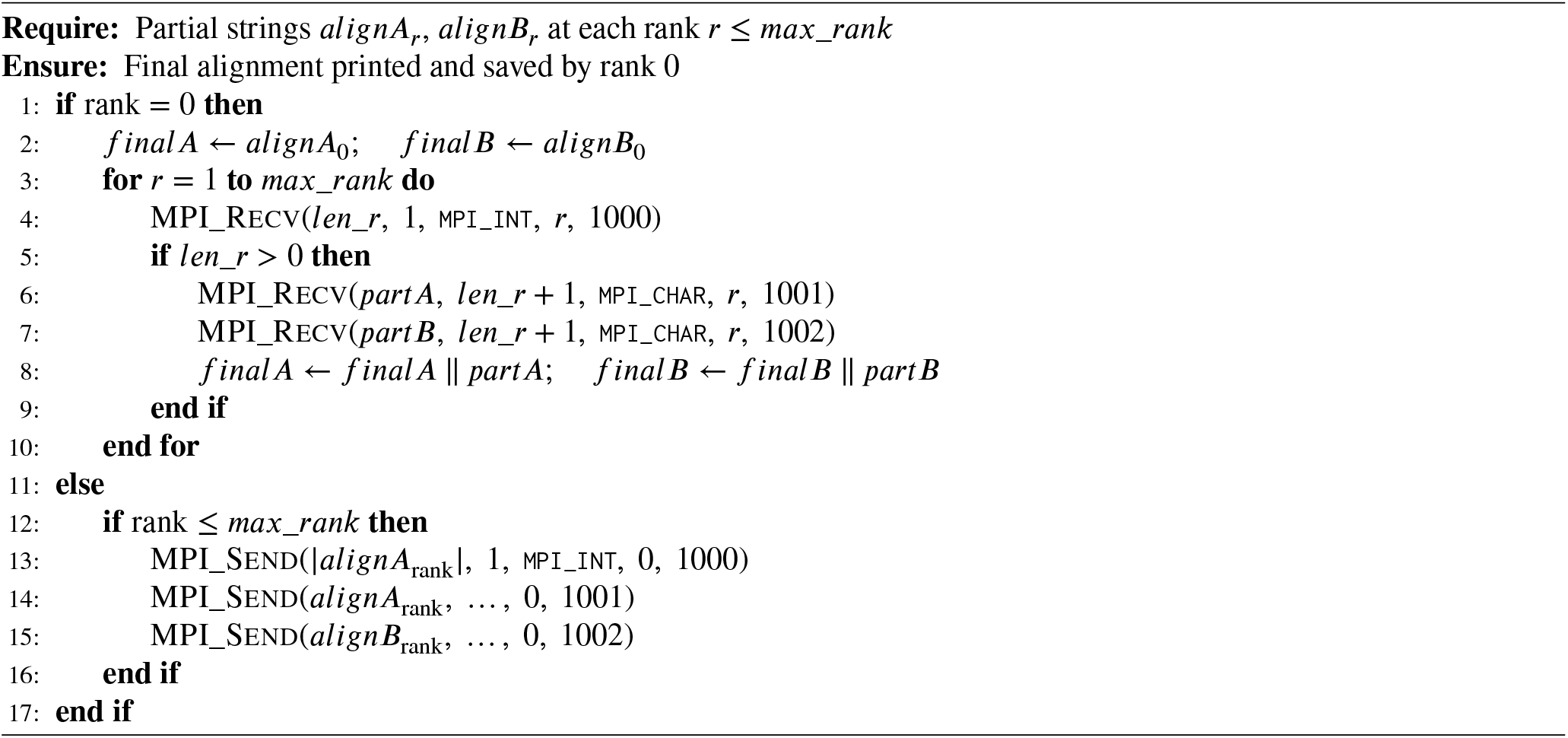

### 5.2. Execution Times Analysis

Table 13 presents the complete execution time data. Sequential time, best MPI time, corresponding processes, and best speedup per dataset are shown in this table. The sequential baseline failed on 50k synthetic, Titin×Titin, and chr1×chr2 real datasets, but the MPI implementation remained successful on all these.

**Table 13.**
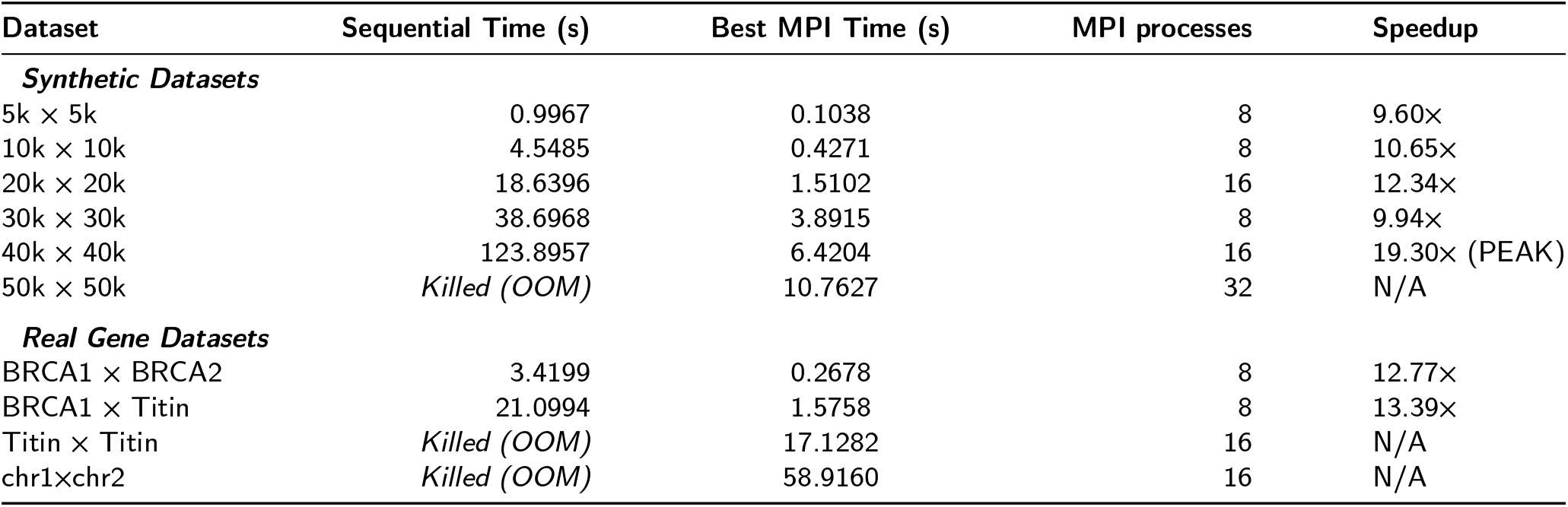
Execution times and peak speedups for Synthetic and Real Gene datasets.

| Dataset | Sequential Time (s) | Best MPI Time (s) | MPI processes | Speedup |
| --- | --- | --- | --- | --- |
| **Synthetic Datasets** |  |  |  |  |
| 5k × 5k | 0.9967 | 0.1038 | 8 | 9.60× |
| 10k × 10k | 4.5485 | 0.4271 | 8 | 10.65× |
| 20k × 20k | 18.6396 | 1.5102 | 16 | 12.34× |
| 30k × 30k | 38.6968 | 3.8915 | 8 | 9.94× |
| 40k × 40k | 123.8957 | 6.4204 | 16 | 19.30× (PEAK) |
| 50k × 50k | *Killed (OOM)* | 10.7627 | 32 | N/A |
| **Real Gene Datasets** |  |  |  |  |
| BRCA1 × BRCA2 | 3.4199 | 0.2678 | 8 | 12.77× |
| BRCA1 × Titin | 21.0994 | 1.5758 | 8 | 13.39× |
| Titin × Titin | *Killed (OOM)* | 17.1282 | 16 | N/A |
| chr1×chr2 | *Killed (OOM)* | 58.9160 | 16 | N/A |

### 5.3. Memory Reduction Analysis

The biggest and clearest achievement of the proposed implementation is the memory reduction, as shown in Table 14. The full score matrix is stored sequentially in the case of the baseline, while the direction matrix is stored as a distributed matrix in the case of our MPI version.

**Table 14.** Memory reduction comparison between Sequential and MPI implementations.

| Dataset | Sequential Memory (MB) | MPI Memory/Rank (MB) | Reduction |
| --- | --- | --- | --- |
| **Synthetic Datasets** |  |  |  |
| 5k × 5k | 190.81 | 3.31 | 57.6× |
| 10k × 10k | 763.09 | 12.57 | 60.7× |
| 20k × 20k | 3052.06 | 24.52 | 124.5× |
| 30k × 30k | 6866.91 | 54.66 | 125.6× |
| 40k × 40k | 12207.64 | 96.72 | 126.2× |
| **Real Gene Datasets** |  |  |  |
| BRCA1 × BRCA2 | 658.99 | 11.07 | 59.5× |
| BRCA1 × Titin | 4521.69 | 75.83 | 59.6× |
| Titin × Titin | Failed (OOM) | 403.86 (16 processes) | N/A |
| chr1×chr2 | Failed (OOM) | 573.06 (16 processes) | N/A |

**Table 15.** Summary of evaluation metrics and peak achievements.

| Metric | Value |
| --- | --- |
| Peak Speedup (Synthetic) | 19.30× (40k×40k, 16 processes) |
| Peak Speedup (Real) | 13.39× (BRCA1×Titin, 8 processes) |
| Largest Dataset Solved | chr1×chr2 (98k×98k) – 58.9160 sec (16 processes) |
| Best Memory Reduction | 126× (40k×40k) |
| Sequential Failures | 50k synthetic, Titin×Titin, chr1×chr2 (OOM) |
| Sweet Spot | 8 processes (best balance of computation vs communication) |

With the 40k×40k dataset, it achieves a 126× memory reduction, where the sequential baseline requires 12.2 GB and MPI requires only 96.72 MB per rank. On chr1×chr2 (98k×98k) real DNA, the MPI program requires 573 MB per rank on 16 processes, whereas the sequential version fails completely.

For weak scaling, the problem size increased in proportion to the number of processes (5k at 1 core, 10k at 2 processes, 20k at 4 processes, 30k at 8 processes, 40k at 16 processes). Figure 9 shows that the execution time remains fairly stable up to 8 processes; at 16 processes, the time increases slightly due to memory pressure and oversubscription on the 8-core host.

**Figure 8.**
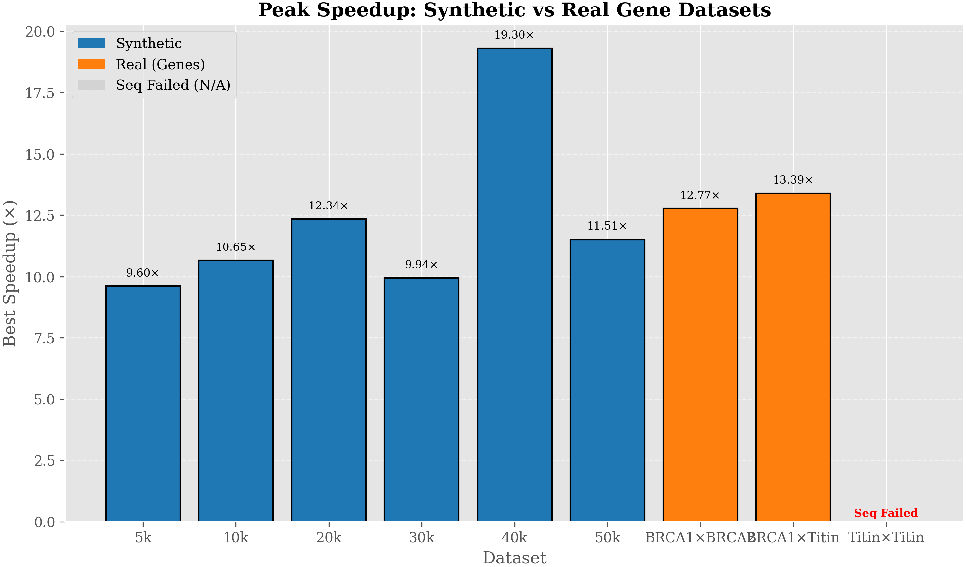
Peak speedup comparison across all tested datasets.

**Figure 9.**
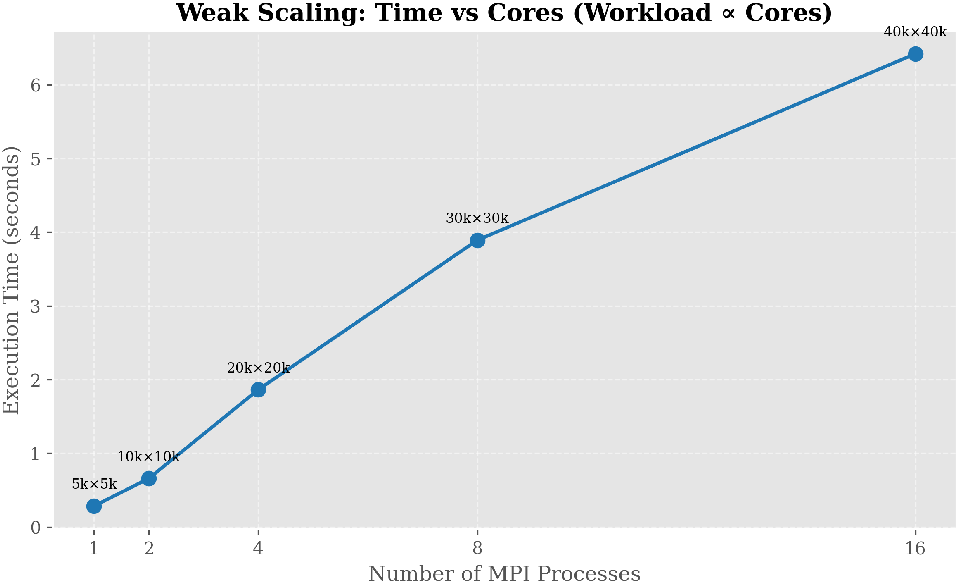
Weak scaling: execution time vs number of MPI processes with workload scaled proportionally to core count..

## 6. Discussion

### 6.1. Scaling Bottlenecks and Limitations

Our MPI implementation has robust scaling behavior but suffers from some bottlenecks that limit the amount of speedup that can be attained. These mainly include pipeline start-up latency; using the row-wise approach, all ranks have to wait for their previous rank’s data before they can start work on their first block, causing the pipeline length to grow with the number of ranks used. This causes scaling to level off at 8 processes and to reduce at 16 processes.

This is compounded by hardware oversubscription; going from 16 to 32 ranks means running more processes than physical cores, and any further increases in the number of ranks will negatively impact the performance. Because all data must fit in distributed memory (7.7 GB into swap), and for even larger datasets, the communication time will become a bottleneck where the ratio between communication and computation will become a scaling bottleneck, as well as the number of `MPI_Send` and `MPI_Recv` calls.

There are also remainder blocks in the data distribution that can cause a load imbalance, where some ranks wait for other ranks to finish. However, as mentioned before, this is acceptable for all but the largest datasets, where the computation dominates the communication, which can be seen from the 19.30× speedup achieved for 40k dataset.

### 6.2. Comparison with Literature and Future Work

We have a balanced implementation compared to previous implementations in the literature. Fitriani et al. [29] report MPI speedups that focus on matrix fill for small test sizes. Nanou [14] requires FPGA hardware, and Bonab et al. [16] require substantially more memory on large gene datasets (746 GB on Titin at 40 cores). Hybrid CPU–GPU approaches [24] remain host-centric for traceback reconstruction.

Instead, we performed SW on a regular workstation/cluster using only 573 MB per rank for the chr1×chr2 dataset, utilizing a distributed Smith-Waterman traceback without gathering the full direction matrix, using rank-to-rank token communication. MPI implementations often have unique communication patterns, but have been shown here to be viable and high-performing methods of computation. We believe this is a strong base for improvements such as an MPI+OpenMP hybrid approach, 2D tiling, and vectorization to reduce scaling bottlenecks.

## 7. Conclusion and Future Work

In this work, we propose a distributed traceback version of the Smith-Waterman algorithm targeted at CPU clusters. We distribute the direction matrix, and perform the traceback in a distributed way, with *O*(*mn*/*P*) memory per node, without gathering the full direction matrix at one node. This method can be used to carry out large-scale sequence alignment, particularly when the conventional sequential baseline fails because of severe memory restrictions.

We experimented on synthetic datasets of sizes ranging from 5k to 50k and obtained good results on real human gene datasets such as BRCA1, BRCA2, and Titin, as well as on chr1×chr2 real DNA chromosome fragments. Our MPI implementation aligns 98k×98k real DNA in 58.9160 seconds on 16 processes, which causes OOM failures in the sequential baseline. From a performance point of view, we obtained a speedup of 19.30× on a 40k×40k synthetic dataset with 16 processes and 13.39× on BRCA1×Titin with 8 processes. In terms of memory usage, the chr1×chr2 real dataset required 573 MB per rank on 16 processes, a significant improvement over the full-matrix approach.

This work addresses a gap left open in prior parallel SW literature, where traceback is often omitted, centralized, or executed serially on a single host or root process. Actor-model implementations can incur very large memory over-heads for manager-based traceback; FPGA implementations provide speedups but rely on specialized hardware with host-side traceback, GPU speedups with traceback remain moderate, and SIMD approaches achieve high throughput without reconstructing alignments. Our work is distinct as a distributed traceback design on CPU MPI clusters keeping the direction matrix partitioned across ranks.

Although successful, the approach has limitations on distributed memory systems. After 8 processes, speedup flattens out because pipeline startup time becomes significant. Hardware oversubscription adds context-switching overhead, and swap-based execution slows large sequential baselines. Communication per pipeline block also limits peak efficiency. These bottlenecks are addressable through 2D tiling, hybrid MPI-OpenMP scaling, and SIMD vectorization.

The most promising approach for future work is a 2D tiling (or wavefront) approach. We currently have a 1D pipeline decomposed row-wise, which means each rank must wait until the previous rank has produced data, resulting in pipeline latency. 2D tiling tiles the matrix across both rows and columns and processes tiles in an anti-diagonal wave-front fashion, enabling higher parallelism within each wave. This could improve scalability to 32, 64, or 128 processes and is compatible with intra-tile OpenMP parallelism.

## CRediT authorship contribution statement

M.F., W.A.: Conceptualization, Data curation, Formal analysis, Investigation, Methodology, Software, Validation, Visualization, Writing - original draft, Writing - review & editing.

W.A.: Supervision, Writing - review & editing, Validation.

## Declaration of Competing Interests

The authors declare that they have no known competing financial interests or personal relationships that could have appeared to influence the work reported in this paper.

## Declaration of Generative AI and AI-assisted Technologies in the manuscript preparation process

During the writing process of this work, the author(s) used DeepSeek to help in the initial drafting of content. After that, content was completely restructured, edited and reviewed thoroughly by the author(s) to maintain scientific accuracy.

## Data Availability

The synthetic datasets used, are generated randomly. Real DNA and chromosomal sequences (BRCA1, BRCA2, etc.) are available on NCBI GenBank website. Upon a reasonable request, MPI source code can be provided by the corresponding author.

## Acknowledgements

The authors would like to thank the Department of Computer Science at the University of Engineering and Technology, Lahore, for providing the computational resources and support necessary to conduct this research.

## Notes

### Competing Interest Statement

The authors have declared no competing interest.

